# When meaning becomes decodable: Linking neural markers of semantic processing and semantic representations

**DOI:** 10.64898/2026.07.16.738961

**Authors:** Gayane Ghazaryan, Aino Saranpää, Tiina Lindh-Knuutila, Marijn van Vliet, Riitta Salmelin

## Abstract

In non-invasive studies of the human brain, semantic processing during language comprehension has been extensively studied using the N400, a component of the electrophysiological evoked response that is strongly modulated by semantic context. More recently, a complementary approach has emerged that uses multivariate pattern analysis to perform neural decoding of semantic vectors from brain activity, operating on the basis that semantic vectors that more closely align with representations in the brain can be more accurately decoded. To consolidate these two approaches, we investigated the relationship between N400 modulation and semantic decoding performance using magnetoencephalography (MEG) in a controlled priming experiment. Twenty-five native speakers of Finnish read word triplets, for which the semantic relatedness between the two primes and the target word was manipulated based on distance in a word2vec embedding space (highly related, moderately related, or unrelated). We found that words presented after unrelated primes elicited higher N400 responses and provided the best examples for training a decoder to map distributed MEG responses to semantic vectors. Semantic information was decodable from approximately 100 to 500 ms after stimulus onset, at all three levels of contextual support. Soon after the N400 peak, neural responses no longer seemed to encode information that could be mapped to context-invariant semantic vectors. This suggests that semantic information is progressively reshaped by linguistic context within and beyond the N400 time window. Semantic decoding therefore offers additional information about representational content that cannot be inferred from the N400 response alone.

## 1. Introduction

When we encounter written text, the brain engages in a complex interplay of perceptual and linguistic processes. Orthographic input, including the shapes and patterns of letters and words, is processed and matched with stored representations in the mental lexicon, enabling word recognition and comprehension (Carreiras et al., 2014; Dehaene et al., 2005). In this way, visual symbols are rapidly transformed into lexical and semantic representations.

This rapid shift from visual word recognition to meaning has traditionally been studied through electrophysiological measures. Among them, the N400 (observed approximately from 300 to 500 ms after stimulus onset) has become the most widely used marker of meaning processing during language comprehension. First described in response to semantically anomalous words in sentences, the N400 was initially associated with semantic incongruity (Kutas and Hillyard, 1980). Subsequent work showed that the N400 amplitude is not specific to semantic incongruity but can be observed for semantically congruent words and systematically modulated by varying their relation to the current semantic context (Kutas and Hillyard, 1984). It can be observed across various types of stimuli including individual words, word lists and non-linguistic stimuli such as pictures (Bentin et al., 1985; Holcomb, 1988; Nigam et al., 1992; Vartiainen et al., 2009; Kutas and Federmeier, 2011; Federmeier, 2022). Expected, related, or repeated items typically show smaller N400 amplitudes, whereas unexpected, weakly related, or poorly supported items show larger amplitudes. These characteristics have made the N400 a central measure of meaning-sensitive processing.

Despite its central role, the functional role of the N400 remains debated (Lau et al., 2008; Kutas and Federmeier, 2011; Federmeier, 2022; Nour Eddine, 2022). Interpretations differ regarding the specific aspect of semantic processing the N400 is thought to reflect. Lexical access accounts argue that the N400 indexes the ease with which lexical-semantic information is accessed from long-term memory (Lau et al., 2008; Kutas and Federmeier, 2011; Federmeier, 2022). The N400 window has further been suggested to index the initial establishment of a stable, detailed semantic representation (Federmeier, 2022). This view draws from evidence for ‘late’ (during the N400 window) semantic processing (Laszlo and Federmeier, 2014; Dufau et al., 2015).

In contrast, post-lexical accounts propose that the N400 reflects processes that occur after the access of the concept, such as integrating it into the preceding context (Brown and Hagoort, 1993; Lau et al., 2008). This account is supported by work identifying ‘early’ (pre-N400 window) semantic processing (Segalowitz and Zheng, 2009; Hauk et al., 2012; Perry, 2022). More recent work has abstained from strict separation between lexical access and integration and has linked the N400 with the degree to which the incoming input updates the current representation state of semantic memory (Nour Eddine, 2022; Nour Eddine et al., 2024).

Although these accounts differ in how they interpret the N400, they converge in treating it as a temporal window of semantic information processing, making it an important reference point for approaches that can elucidate when and how semantic information is represented in neural activity. One such approach relies on multivariate pattern analysis and neural decoding. Rather than focusing on response magnitude at selected sensors or time windows, decoding approaches ask what information can be recovered from distributed patterns of brain activity (Haynes and Rees, 2006; Haxby et al., 2014). In language research, this enables us to examine whether neural responses contain information that maps onto individual word meanings and the semantic relationships between them.

This approach is particularly powerful when paired with computational semantic models, which represent words in multidimensional spaces that allow semantic relationships between words to be quantified (Landauer and Dumais, 1997; Mikolov et al., 2013; Huth et al., 2016). Word2vec (Mikolov et al., 2013) is one such model. It represents words as vectors based on co-occurrence statistics from a large corpus. These vectors can be considered context-invariant estimates of meaning. Words with similar co-occurrence patterns are represented by similar vectors, and semantic relatedness between two words can be quantified by the distance between the corresponding vectors. Several neural decoding studies have shown that distributed brain responses can be mapped onto word2vec embeddings (Ghazaryan et al., 2023b,a; Hultén et al., 2021; Sassenhagen and Fiebach, 2020), and words with lower word2vec distance to a preceding word or context generally elicit smaller N400 responses than those with greater distance (Ettinger et al., 2016; Dudschig et al., 2026).

Written words, while commonly used in studies of the N400 response, are a challenging case for neural decoding. Unlike pictures, written words do not visually depict the meanings they convey. Successful decoding therefore requires identifying semantic information in the neural response that reflects lexical and conceptual processing and is not directly tied to the visual form of the stimulus (Dehaene et al., 2005; Binder et al., 2009; Pulvermüller, 2013). Decoding semantic information from time-sensitive neuroimaging recordings (e.g. EEG and MEG) to written words has therefore remained relatively uncommon and technically challenging (Chan et al., 2011; Rybář and Daly, 2022; Ghazaryan et al., 2023b). Recent studies have shown that such decoding is feasible under suitable experimental conditions (Hultén et al., 2021; Hakala et al., 2024; d’Ascoli et al., 2025), opening the possibility of combining classical univariate analyses and multivariate methods within the same study and examining the link between the approaches (Wang and Kuperberg, 2024). A specific intriguing question is whether contextual modulation of the N400 as captured by univariate evoked analyses is associated with differences in how reliably word meanings can be decoded from multivariate patterns.

In the present study, we tested this relationship using MEG responses to written words in a controlled semantic priming paradigm. Target words appeared at the end of word triplets and were preceded by two primes whose semantic relatedness to the target was systematically varied. This design created graded contextual support for the target while preserving a repeated target word structure suitable for decoding. We combined conventional analyses of evoked responses in the N400 window with semantic decoding to ask whether conditions that attenuate or enhance the N400 also differ in how reliably target-word meaning can be recovered from neural activity. We further examined the timing of semantic decodability relative to the N400 window, examining both the pre-N400 and post-N400 windows, and the generalization of information between levels of contextual support and through time. By linking N400 modulation with semantic decoding, this study provides an empirical bridge between univariate and multivariate approaches to the analysis of semantic processing in the brain.

## 2. Methods

### 2.1. Participants

The study involved a group of 26 right-handed native Finnish speakers (females/males 14/12), aged between 19 and 45, with a mean age of 24.77 (SD: 7.16). Participants had normal or corrected-to-normal vision. The study was approved by the Research Ethics Committee of Aalto University and the participants provided informed consent before participating. Data from one participant were omitted from the analysis due to technical problems with the MEG recordings, resulting in a final sample of 25.

### 2.2. Stimuli

We generated word triplets where the final word (target) was semantically related to the two preceding words to varying degrees. The words were presented to the participants in the order of Prime 1, Prime 2 and Target during the experiment. However, in constructing these triplets, we work backwards, from the targets to the primes, using a pre-trained Finnish word2vec model trained on Finnish Internet Parsebank corpus (Luotolahti et al., 2015; Mikolov et al., 2013; Kanerva et al., 2014). This approach allowed us to systematically manipulate the semantic relatedness between targets and primes.

We selected 70 target nouns, from 14 categories: food, animals, tools, body parts, clothing, vehicles, plants, culture, time, emotions, economy, beliefs, characteristics, and roles. For each target word, we then computed distances to 15000 common Finnish nouns in the word2vec embedding space. This resulted in a semantic distance profile for each target word relative to the most frequent nouns in the language. Recognizing that words vary in semantic neighborhood density within the word2vec embedding space, we employed a relative distance metric unique to each target word. For each target word, we sorted the common nouns based on their cosine distance to that target and divided them into groups according to specific percentile ranges to represent different levels of semantic relatedness:

- High Relatedness: 0.25% closest words to the target.
- Moderate Relatedness: 0.5 to 1% closest words to the target.
- Unrelated: further than the closest 10% of words.

This approach accounted for variations in semantic density, meaning that a specific distance value could represent moderate relatedness for one target but high relatedness for another. By using percentile-based groupings rather than fixed distance thresholds, we ensured consistent levels of relatedness across all target words, appropriately scaling the levels of relatedness relative to each target’s neighborhood.

For each Target, we first generated three sets of Prime 2 words for each relatedness group. This allowed us to manipulate the semantic relatedness between Prime 2 and the target word across three distinct levels for every target. Subsequently, for each Prime 2, we identified a Prime 1 that was highly semantically related to Prime 2. Specifically, Prime 1 was chosen from the top 0.25% closest words to Prime 2, ensuring a strong semantic connection between the two primes and establishing a robust contextual foundation. This methodology resulted in the creation of word triplets for each target word, where (1) prime 1 and prime 2 were highly related, providing a strong contextual cue and (2) prime 2 and the target word had systematically manipulated levels of semantic relatedness. Prime 1 (*F* (2, 768) = 0.32, *p* = 0.72) and prime 2 (*F* (2, 768) = 1.62, *p* = 0.2) words did not differ in length across the levels of semantic relatedness. There were also no differences in associated frequency, (*F* (2, 768) = 0.02, *p* = 0.98) and (*F* (2, 768) = 0.74, *p* = 0.48), respectively. Supplementary figures S1, S2, S3 and S4 show the word frequencies and lengths for the prime words for each target.

From these generated trials, we selected four trials per target in each level of relatedness. Thus, we had a total of 840 unique triplets. Each triplet was presented only once to participants, ensuring that responses were based on unique, non-repeating stimuli. Figure 1 shows the distribution of distances between primes and targets for the final set of stimuli.

**Figure 1:**
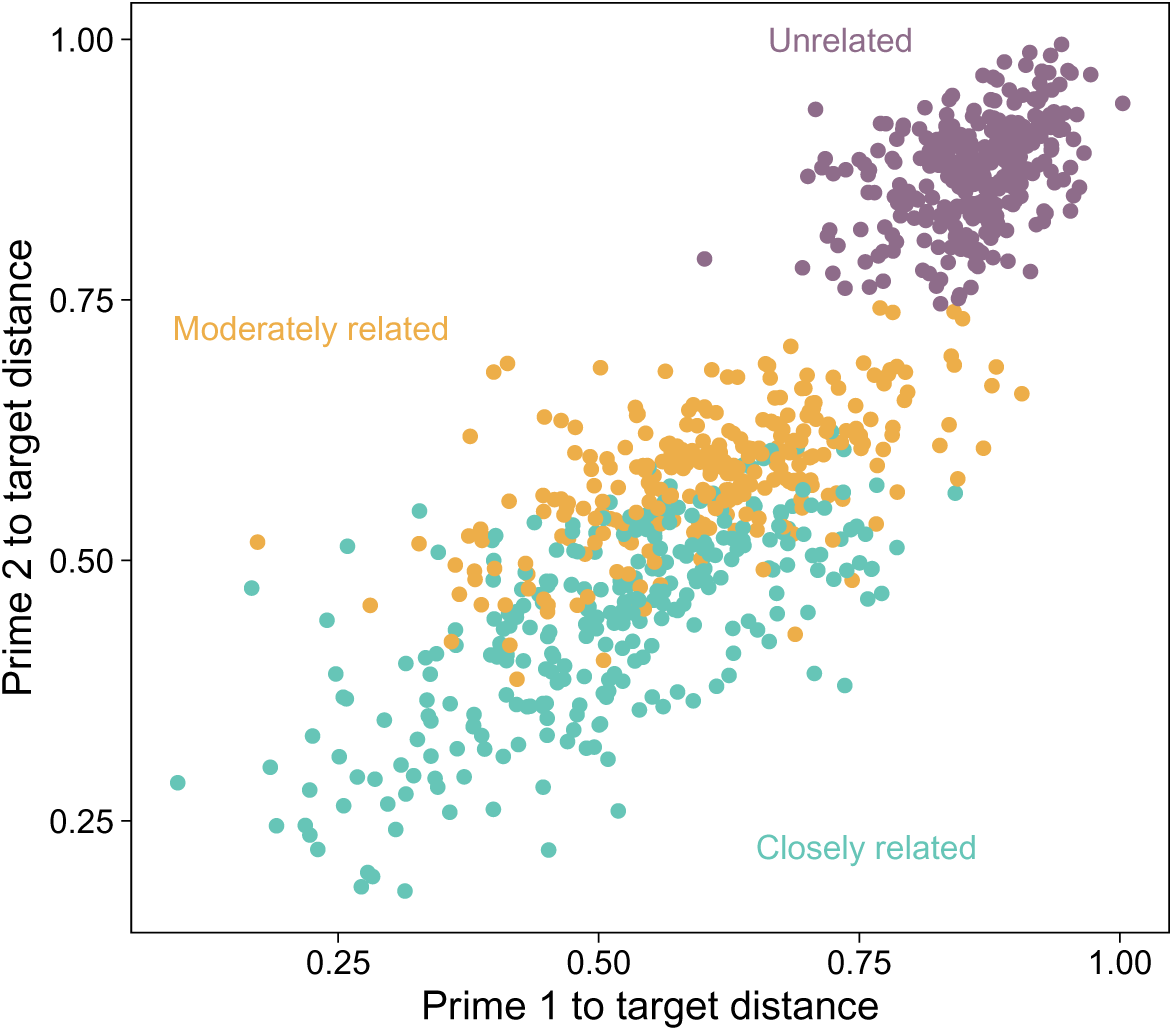
Distributions of cosine distances between prime and target word2vec vectors for the three levels of relatedness.

### 2.3. Procedure

Participants were seated comfortably at a reclined position (60 degrees) in a dimly lit magnetically shielded room, positioned approximately 120 cm from the display screen. Stimuli were presented using Presentation software (Neurobehavioral Systems, Inc.) by a projector with a refresh rate of 120 Hz.

Each trial (Figure 2A) began with a fixation cross displayed at the center of the screen for 400 ms to orient the participant’s attention. Following the fixation, the three words were presented sequentially: prime 1, prime 2, target word. Each word appeared individually at the center of the screen for 400 ms. Following the presentation of each prime, a blank screen was displayed for 400 ms, and a blank screen of randomly determined interval ranging from 800 to 1200 ms was presented after the target word to prevent anticipation of the next trial’s onset.

**Figure 2:**
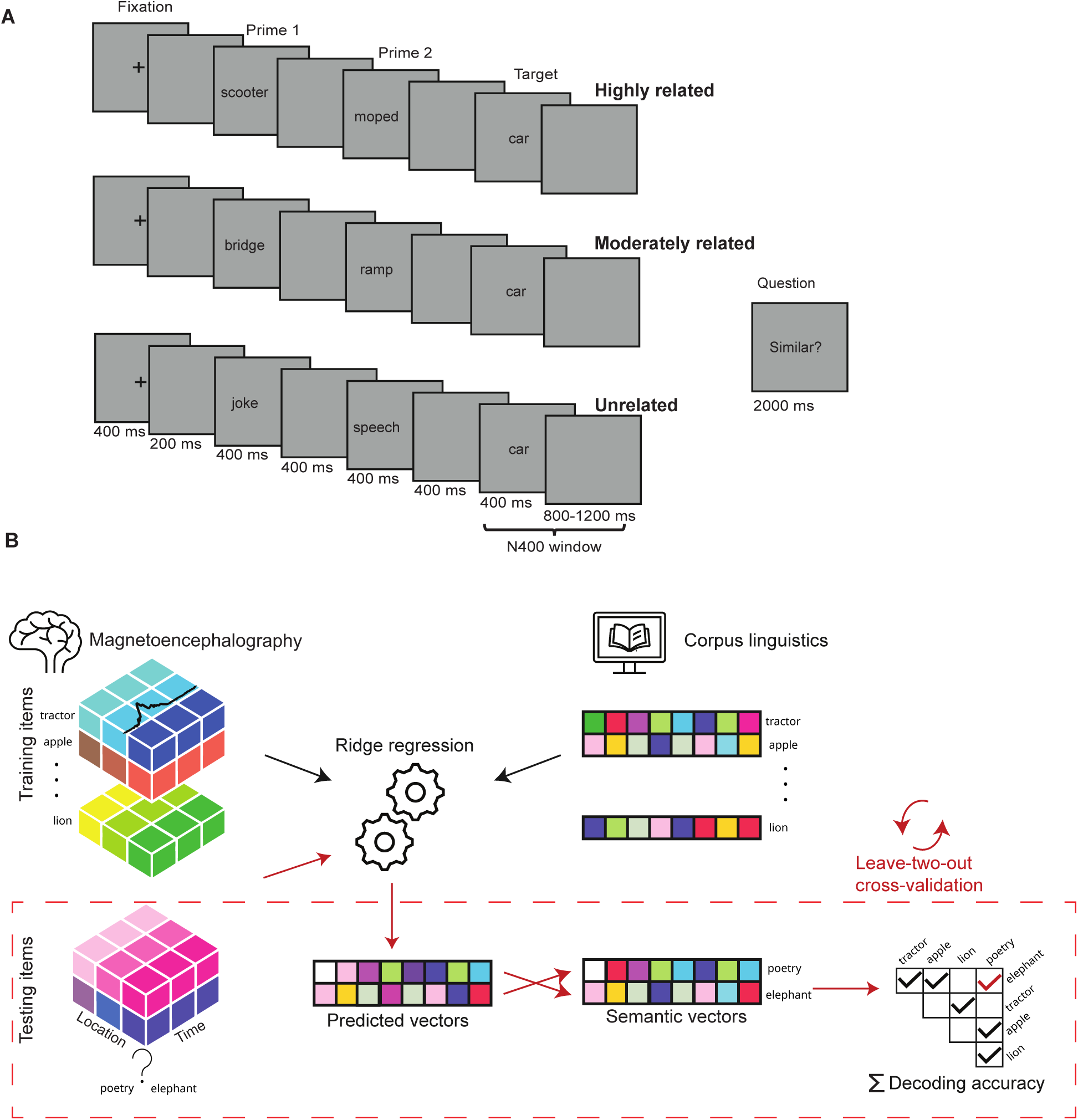
A. Example trial from the experiment showing timings of stimuli presentation. 10% of the trials were followed by a yes/no question to ensure participant engagement. B. Overview of neural decoding approach used to map brain signals to concepts.

The experiment consisted of 4 blocks, each containing 210 unique trials. In each block, 70 target words were presented once in each of the three relatedness levels. Across the entire experiment, each target word was presented four times in each relatedness level. Targets were always preceded by primes that had not appeared before with the same target. All the blocks were presented in one measurement session, and the order of the blocks was pseudorandomized across participants to mitigate sequence effects.

Participants were instructed to determine whether the words in the presented triplet were related to each other. To ensure task engagement while minimizing artifacts associated with motor responses, participants were required to make an explicit response after only 10% of the trials, randomly assigned as catch trials. After these catch trials, participants indicated whether the words were similar to each other by lifting their index fingers on optical response pads, using different hands for each response. The response mapping (i.e., which hands corresponded to ‘yes’ and ‘no’) was switched after two blocks to control for any lateralized motor activity. Participants were given a rest period after two blocks. The mean proportion of ‘yes’ responses was 85.7% for highly related words (SD = 10.0%), 66.4% for moderately related words (SD = 16.8%), and 10.5% for unrelated words (SD = 6.57%).

### 2.4. Data acquisition

Magnetoencephalography (MEG) data were collected using a 306-channel whole-head MEG system (MEGIN TRIUX neo, MEGIN Oy, Espoo, Finland) at the Aalto NeuroImaging MEG Core facility. The system consists of 204 planar gradiometers and 102 magnetometers. The data analysis was performed on the 204 planar gradiometers. Participants’ head positions were continuously monitored throughout the experiment using five head position indicator (HPI) coils placed at known locations relative to identifiable anatomical landmarks (nasion and preauricular points).

To monitor ocular artifacts, eye movements and blinks were recorded using two pairs of electrodes. One pair was positioned above and below the left eye to detect vertical eye movements, while the other pair was placed at the outer corners of the eyes to capture horizontal movements. MEG signals were lowpass-filtered online at 330 Hz and data were sampled at a rate of 1000 Hz.

Anatomical magnetic resonance imaging (MRI) scans were obtained for each participant using a Siemens Magnetom Skyra 3.0 T MRI scanner at the Aalto NeuroImaging Advanced Magnetic Imaging (AMI) Centre. T1-weighted MP-RAGE and T2-weighted SPC SAG sequences were acquired to facilitate accurate co-registration of MEG and MRI data.

### 2.5. Data preprocessing

MEG data preprocessing was performed using the MNE-Python software package (Gramfort, 2013). The initial step involved visually examining the MEG data to detect and mark any channels with poor data quality, i.e. with high noise or no reliable signal. To suppress external noise sources, we utilized spatiotemporal signal space separation (tSSS; Taulu and Simola, 2006)). Data from different blocks for each participant were brought to the same head position to maintain consistency across recordings.

We then applied a band-pass filter between 0.1 and 40 Hz to the continuous MEG data to reduce low-frequency drifts and high-frequency noise. To mitigate artifacts caused by heartbeats and eye movements, independent component analysis (ICA) was conducted. For ICA, we used continuous data that were high-pass filtered at 1 Hz to reduce the impact of slow drifts on the decomposition (Jas et al., 2018). Components associated with cardiac and ocular artifacts were visually identified and removed from the continuous data high-pass filtered at 0.1 Hz.

We segmented the data into epochs of 1100 ms, including a 100-ms pre-stimulus baseline. Epochs corresponding to the same target words were then averaged. In order to minimize positional differences across participants, head positions of individuals were transformed to a common reference position. Following alignment, data from each participant were averaged to obtain grand-averaged evoked responses, calculated separately for each target word within each level of relatedness. We then extracted the time window from 0 to 1000 ms post-stimulus and resampled the data into 20-ms time bins. Thus we arrived at a data matrix of 70 targets× 204 gradiometer channels × 50 time points × 3 relatedness levels.

### 2.6. Zero-shot decoding

To explore how brain responses correspond to semantic features, we utilized a zero-shot decoding approach with multivariate ridge regression. This is an established approach used in previous work (e.g., Ghazaryan et al., 2023b; Kivisaari et al., 2019; Hultén et al., 2021) and is used to evaluate the ability to decode new examples that models have not been trained on (Xian et al., 2019). Before decoding analysis, the data were reshaped so that each timepoint-sensor combination served as an individual predictor in the regression model. We then transformed the brain response data by standardizing each timepoint-sensor combination to have a mean of 0 and a standard deviation of 1.

We then fit the regression models and assessed the decoding accuracy using a leave-two-out scheme, commonly used to evaluate zero-shot decoding (Mitchell et al., 2008; Fyshe et al., 2019; Kivisaari et al., 2019; Ghazaryan et al., 2023b). In each iteration, two target items were left out from the training set and the model was trained on the remaining data and then used to predict the semantic vectors from the brain responses of the two excluded target words. Decoding was considered successful if the predicted vector for each target word was closer in semantic space to its own true vector representation. Thus, predicted vectors for both left-out items needed to be more similar to their true semantic representations than to the other for the decoding to be deemed successful. This procedure was exhaustively repeated for all possible pairs of left-out items, and we ensured valid cross-validation by standardizing the brain responses within each cross-validation fold, excluding the left-out targets.

Such decoding analyses were performed for each relatedness level separately. Additionally, a pooled model was used which was trained on all relatedness levels and tested on each separately. In this case, to maintain the zero-shot nature of the testing, all instances of the test items were removed from the training sets. We investigated the decoding accuracy of models trained and tested at separate time windows: 100–300 ms, 300–500 ms and 500–700 ms.

### 2.7. Cross-temporal and cross-relatedness level decoding

To capture the temporal dynamics of semantic processing, we performed time-resolved decoding using a sliding window approach. Each window spanned 100 ms and overlapped with the next by 80 ms. This method allowed us to investigate how the brain’s representation of semantic features evolves over time. For each time window, the zero-shot decoding approach described above was employed, resulting in decoding accuracy over time.

To evaluate the generalizability of semantic representations across different times (King and Dehaene, 2014) and levels of semantic relatedness, we extended our analysis to include cross-temporal and cross-relatedness level decoding. In the cross-temporal decoding, the model was trained on data from one 100 ms time window and tested on data from a different 100 ms window. This tested the model’s ability to generalize learned semantic representations to a new temporal context. We performed this analysis both within the same level of semantic relatedness and across different levels. For cross-relatedness level decoding the model was trained on data from one relatedness level and tested on data from another level. By combining these approaches, we aimed to gain a comprehensive understanding of how semantic information is encoded and generalized in the brain over time and across different levels of contextual support.

### 2.8. Cortical-level analysis

Each participant’s cortical surface was reconstructed from their anatomical MRI scans using the FreeSurfer software package (Fischl, 2012). Source-level estimates of neural responses for each concept were computed using the minimum norm estimation (MNE) method. A single-layer boundary element model (BEM) was employed for forward computation. In calculating the inverse solution, a loose orientation constraint of 0.2 and a depth weighting parameter of 0.8 were applied. An empirical noise covariance matrix was estimated from the 100-ms pre-stimulus intervals across all targets. Dynamical Statistical Parametric Maps (dSPMs) were calculated over the entire cortical area. Individual source estimates were morphed onto the standard FreeSurfer template brain (fsaverage). Contrasts between relatedness levels were first computed for each participant and then assessed at the group level using spatio-temporal cluster-based permutation tests.

We additionally performed zero-shot decoding using source-localized data. This builds on previous work (Magnabosco and Hauk, 2024; Keitel et al., 2020; Wu et al., 2024), where source-space decoding of target words was performed, albeit not in a zero-shot framework. For this analysis, source estimates morphed onto the standard template brain were first averaged across participants. We then used a full-brain searchlight approach (Kriegeskorte et al., 2006) in the three time windows of interest (100–300 ms, 300–500 ms, 500–700 ms). Models were trained and tested using data from vertices within patches of radius 20 mm.

### 2.9. Statistics

To establish statistical significance for our window-based decoding results, we conducted permutation tests. We disrupted the mapping between brain data and semantic vectors by shuffling the data, effectively breaking any true correspondence between the brain responses and semantic vectors. This process was repeated 1,000 times, and the zero-shot decoding with the leave-two-out scheme was performed on each permuted dataset (following Valente et al., 2021). This resulted in a null distribution, which was used to determine the significance of the decoding accuracy of models trained and tested on the true data, and the differences between relatedness levels and time windows. For the temporal generalization maps, we used two-dimensional time x time cluster-based permutation tests (Maris, 2012), similarly to Dijkstra et al. (2018) and Hebart et al. (2018). For source-space searchlight decoding we used spatial cluster-based permutation tests, similar to Keitel et al. (2020). Cluster-inclusion thresholds for both analysis types were *p* < .05. Cluster permutation testing was performed using the permuco and permuco4brain R packages (Frossard and Renaud, 2021). Multiple comparisons were controlled for using the False Discovery Rate correction (Benjamini and Hochberg, 1995).

## 3. Results

### 3.1. Semantic relatedness modulates the evoked response

Semantic relatedness modulated the evoked response to the target item in the N400 window (300–500 ms). Mean amplitudes for each target item over this interval were analyzed for three MEG sensor groups associated with areas known to exhibit N400 effects. Repeated-measures ANOVAs with relatedness level as a within-item factor indicated main effects of relatedness level in the left frontal sensors (*F* (1.92, 132.52) = 15.31, *p <* 0.001), left parietal sensors (*F* (1.95, 134.78) = 4.38, *p* = 0.015), and left temporal sensors (*F* (1.87, 128.93) = 24.87, *p* < .001; Figure 3).

**Figure 3:**
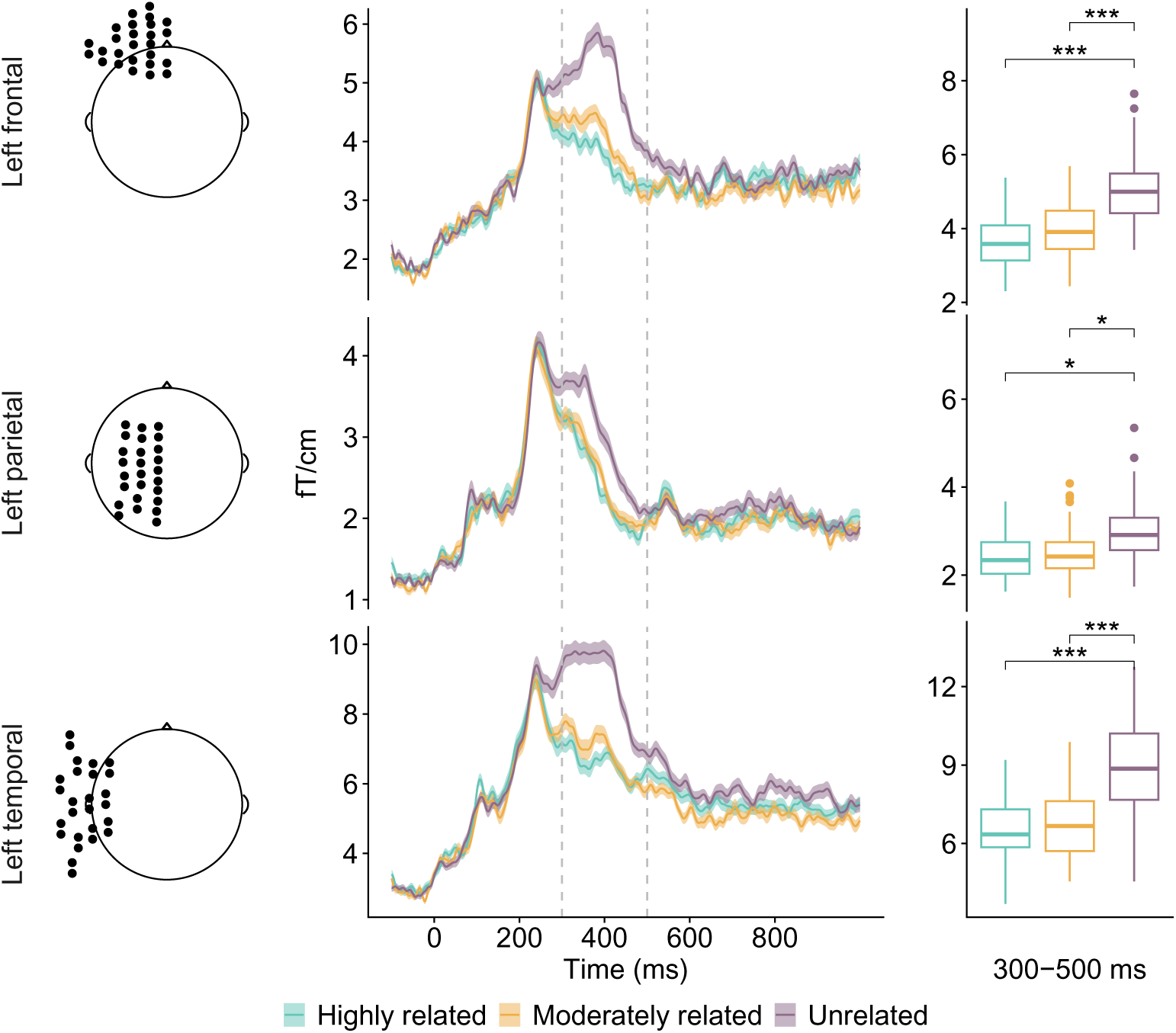
Left: MEG sensor layouts of the regions of interest. Middle: Evoked responses for each relatedness level averaged over items. Right: Differences in signal amplitude of target items depending on relatedness level in the N400 window (300–500 ms). Mean amplitudes for the unrelated, moderately related, and highly related contexts were 5.05 ± 0.94, 3.96 ± 0.73, and 3.68 ± 0.68 fT/cm in the left frontal sensors; 2.96 ± 0.64, 2.49 ± 0.55, and 2.39 ± 0.47 fT/cm in the left parietal sensors; and 8.92 ± 1.70, 6.83 ± 1.27, and 6.56 ± 1.10 fT/cm in the left temporal sensors. ∗ = *p* < .05, ∗ ∗ ∗ = *p* < .001.

Pairwise contrasts, corrected using FDR across the nine pairwise tests, showed the same pattern in all three sensor groups: amplitudes were higher for unrelated targets than either the moderately related or highly related targets, whereas the highly related and moderately related targets did not differ significantly. The unrelated targets induced significantly higher amplitudes than the others in all three sensor groups (left frontal: both *p <* 0.001; left parietal: *p* = 0.020 and 0.031; left temporal: both *p <* 0.001, whereas the highly related and moderately related targets did not differ (left frontal: *p* = 0.736; left parietal: *p* = 0.650; left temporal: *p* = 0.082. This modulation was mirrored at the cortical level. There was a significant difference between the unrelated and moderately related targets (cluster *p* = 0.002) and between the unrelated and highly related targets (cluster *p* = 0.002; Figure 4). Again, no significant difference was detected between the highly related and moderately related targets (cluster *p* = 0.756).

**Figure 4:**
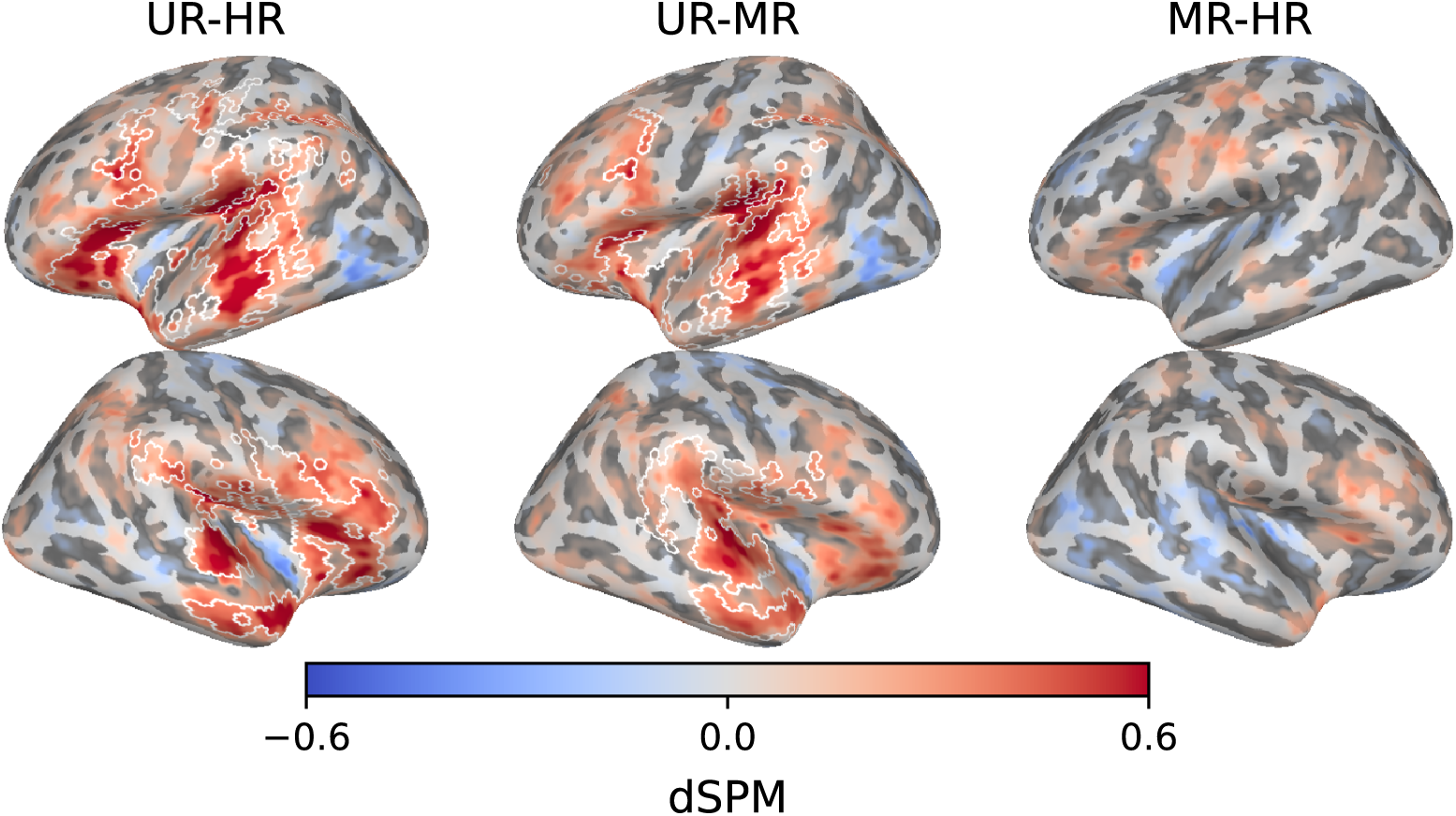
Source-localized activation differences across relatedness levels in the N400 window (300–500 ms). Mean dSPM across participants is shown. White borders indicate spatial extent of spatio-temporal clusters with associated p-values lower than .05. HR = highly related, MR = moderately related, UR = unrelated.

### 3.2. Semantic decoding is successful around the N400 window

Target words were decodable from the MEG signal in a zero-shot framework, indicating that a generalizable mapping from MEG signal to semantic space was learned (Figure 4), left). All reported p-values were derived from permutation tests and corrected for multiple comparisons using FDR across 21 tests: nine tests against chance, six within-context comparisons between time windows, and six between-context comparisons within time windows.

In the pre-N400 window (100–300 ms), decoding was significantly above chance for highly related targets (62.6%, *p* = 0.006) and unrelated targets (59.5%, *p* = 0.028, but not for moderately related targets (44.3%, *p* = 0.897). Between-relatedness level comparisons showed the same pattern: decoding accuracy was higher for both highly related and unrelated targets than for moderately related targets (highly related vs. moderately related: *p* = 0.0047; unrelated vs. moderately related: *p* = 0.0026).

In the N400 window (300–500 ms), decoding accuracy was significantly above chance for unrelated targets (83.0%, *p* = 0.002) and moderately related targets (69.3%, *p* = 0.002), but not for highly related targets (57.9%, *p* = 0.053). Between-level comparisons confirmed this ordering: decoding accuracy was higher for unrelated than moderately related targets (*p* = 0.006), and higher for moderately related than highly related targets (*p* = 0.028)

In the post-N400 window (500–700 ms), decoding accuracy decreased relative to the N400 window across all three relatedness levels. Unrelated targets remained significantly above chance (64.9%, *p* = 0.0023), whereas highly related targets (48.8%, *p* = 0.635) and moderately related targets (52.5%, *p* = 0.317) did not. Decoding accuracy was higher for unrelated than moderately related targets (*p* = 0.0105), while highly related and moderately related targets did not differ significantly (*p* = 0.276). Thus, after the N400 window, target-word decodability was reduced overall and remained reliable only for unrelated targets.

Within-relatedness level comparisons confirmed that decoding peaked in the N400 window for both moderately related and unrelated targets. In both cases, N400-window accuracy was significantly higher than accuracy in the pre-N400 and post-N400 windows (all *p* = 0.003). Highly related targets showed no reliable decrease from the pre-N400 to the N400 window (*p* = 0.173).

### 3.3. Decoding models generalize through time and across relatedness levels

Temporal generalization maps were computed for each train–test combination of relatedness level and evaluated using cluster-based permutation tests (Figure 6, left three columns). Among the within-context combinations, there was significant temporal generalization only for unrelated-unrelated (*p* = 0.020), whereas highly related–highly related did not survive FDR correction (*p* = 0.054) and moderately related–moderately related was not significant. Temporal generalization was nevertheless observed for all cross-context train–test combinations (all *p ≤* 0.036). Across the combinations showing temporal generalization, this pattern was most evident in the pre-N400 and N400 portions of the maps. Together, these results indicate that the decodable semantic features remained sufficiently stable over time and were shared across relatedness levels.

### 3.4. Pooled training strengthens semantic decoding across contexts

Given the indication of generalization across relatedness levels, training data were pooled across all three levels and decoding analysis was repeated. We continued to test on data from each relatedness level separately but retained the zero-shot framework by excluding all instances of the test items from the pooled training set. As shown in Figure 5 (right), with pooled training, decoding accuracy was significantly better than chance in the pre-N400 window when test items were from the highly related (76.8%, *p* = 0.001), moderately related (67.6%, *p* = 0.001), or unrelated (73.0%, *p* = 0.001) contexts. In the N400 window, decoding accuracy was again significantly higher than chance for all three contexts (highly related: 73.3%, moderately related: 71.3%, unrelated: 79.9%; all *p* = 0.001), and did not differ from the pre-N400 window for any context (all *p >* 0.1). In the post-N400 window, decoding accuracy declined relative to the N400 window in all three contexts (all *p* = 0.002), but remained significantly above chance for the unrelated context (62.4%, *p* = 0.004). Decoding accuracy in the highly related (48.9%, *p* = 0.586) and moderately related (50.1%, *p* = 0.514) contexts was not significantly above chance. Between-context comparisons showed no reliable differences in the pre-N400 or N400 windows (all *p >* 0.05), whereas in the post-N400 window decoding accuracy was higher in the unrelated than the moderately related context (*p* = 0.0248). Temporal generalization under pooled training was also observed for all three test contexts (all *p* = 0.020); Figure 6, right column), again primarily seen in the pre-N400 and N400 windows. Together, these results indicate that pooling training data improved estimation of the relationship between MEG patterns and semantic representations.

**Figure 5:**
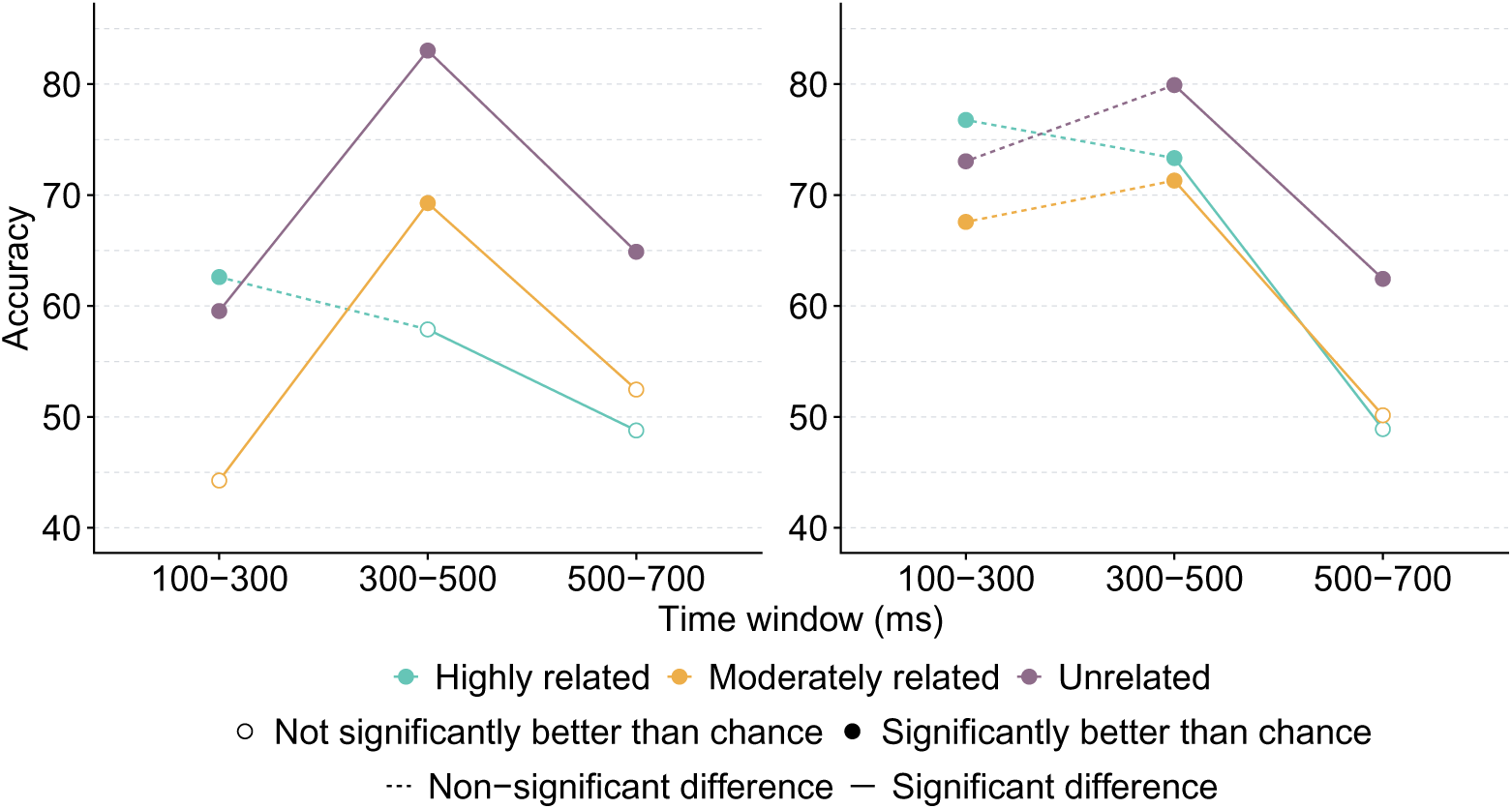
Zero-shot decoding accuracy of target words from grand average MEG data. Left: Results for models trained and tested on data from the same relatedness levels. Right: Results for models trained on data pooled across relatedness levels and tested on individual contexts.

### 3.5. Cortical-level searchlight decoding highlights regions of semantic processing

Cortical-level searchlight decoding maps were computed using pooled training data for each relatedness level and time window (Figure 7). Cluster-based permutation tests showed that decoding accuracy was significantly above chance in all windows for all three relatedness levels (all *p <* 0.012). Cluster-based permutation tests cannot pinpoint specific regions in which decoding was significantly better than chance (Sassenhagen and Draschkow, 2019). However, the cortical areas which tended toward higher decoding accuracy broadly align with areas associated with semantic processing, such as temporal, inferior parietal, and inferior frontal cortex (Lau et al., 2008; Binder et al., 2009; Lambon Ralph et al., 2017).

## 4. Discussion

The present study brought together two widely used but largely separate approaches to study semantic processing: the N400, a well-established neural marker of semantic processing, and multivariate decoding of semantic representations. Using a controlled word triplet paradigm, we varied the degree of semantic relatedness from two prime words to a target word. The same target words were presented repeatedly across relatedness levels, but always in unique triplets, creating a design suitable for both conventional evoked-response analysis and more recent zero-shot semantic decoding. This allowed us to examine whether conditions that modulate the N400 also change how reliably target word meaning can be recovered from distributed MEG activity and at what time the target meaning is present in the brain.

Semantic relatedness modulated the evoked response in the expected direction. Targets preceded by unrelated primes produced larger N400 responses than targets preceded by moderately or highly related primes. This pattern is consistent with extensive evidence that N400 amplitude is larger when incoming meaningful input is weakly supported by preceding semantic information, and smaller when the target is expected or related (Kutas and Hillyard, 1980; Lau et al., 2008; Kutas and Federmeier, 2011; Federmeier, 2022). The effect was observed in left frontal, parietal, and temporal MEG sensor groups and was mirrored in cortical-level estimates, indicating that the manipulation reliably distinguished unrelated targets from targets with greater contextual support.

Semantic information about target words was decodable from the whole-brain MEG signal. In the zero-shot decoding framework, models learned a mapping between MEG activity and context-invariant semantic vectors and then applied that mapping to unseen target words. Successful decoding therefore indicated that the MEG signal contained information that related to semantic relationships among target words. This approach builds on previous work relating semantic embeddings and neural activity (Mitchell et al., 2008; Hultén et al., 2021; Ghazaryan et al., 2023a).

Decoding accuracy in the N400 window appeared to vary with the N400 amplitude: unrelated targets elicited the largest N400 responses and showed the strongest decodability, followed by moderately related and then highly related targets (Figure 5, left). However, this was not the full picture. The pooled-training analysis showed that when models were trained across all three relatedness levels, target words were reliably and similarly decodable irrespective of contextual support, from both the pre-N400 and N400 windows (Figure 5, right). Temporal generalization further indicated that the decodable information was relatively stable in these windows (Figure 6), and cortical-level searchlight decoding highlighted broadly similar areas for both the pre-N400 and N400 windows (Figure 7) overlapping with cortical areas implicated in semantic processing, including temporal, inferior parietal, and inferior frontal regions (Binder et al., 2009; Huth et al., 2016; Lambon Ralph et al., 2017). Together, these findings suggest that MEG responses to written words from approximately 100 to 500 ms after stimulus onset contain relatively stable target-specific semantic information that can be mapped onto context-invariant embeddings. This result does not strictly refute accounts emphasizing ‘early’ (pre-N400 window) semantics (Segalowitz and Zheng, 2009; Sereno et al., 2020; Perry, 2022; Hauk et al., 2012) nor ‘late’ (N400 window) semantics (Laszlo and Federmeier, 2014; Dufau et al., 2015), or the associated interpretations of the N400. However, the results do suggest that stable representations appear already in the pre-N400 window, indicating that lexical-semantic access likely occurs earlier than the N400 peak.

**Figure 6:**
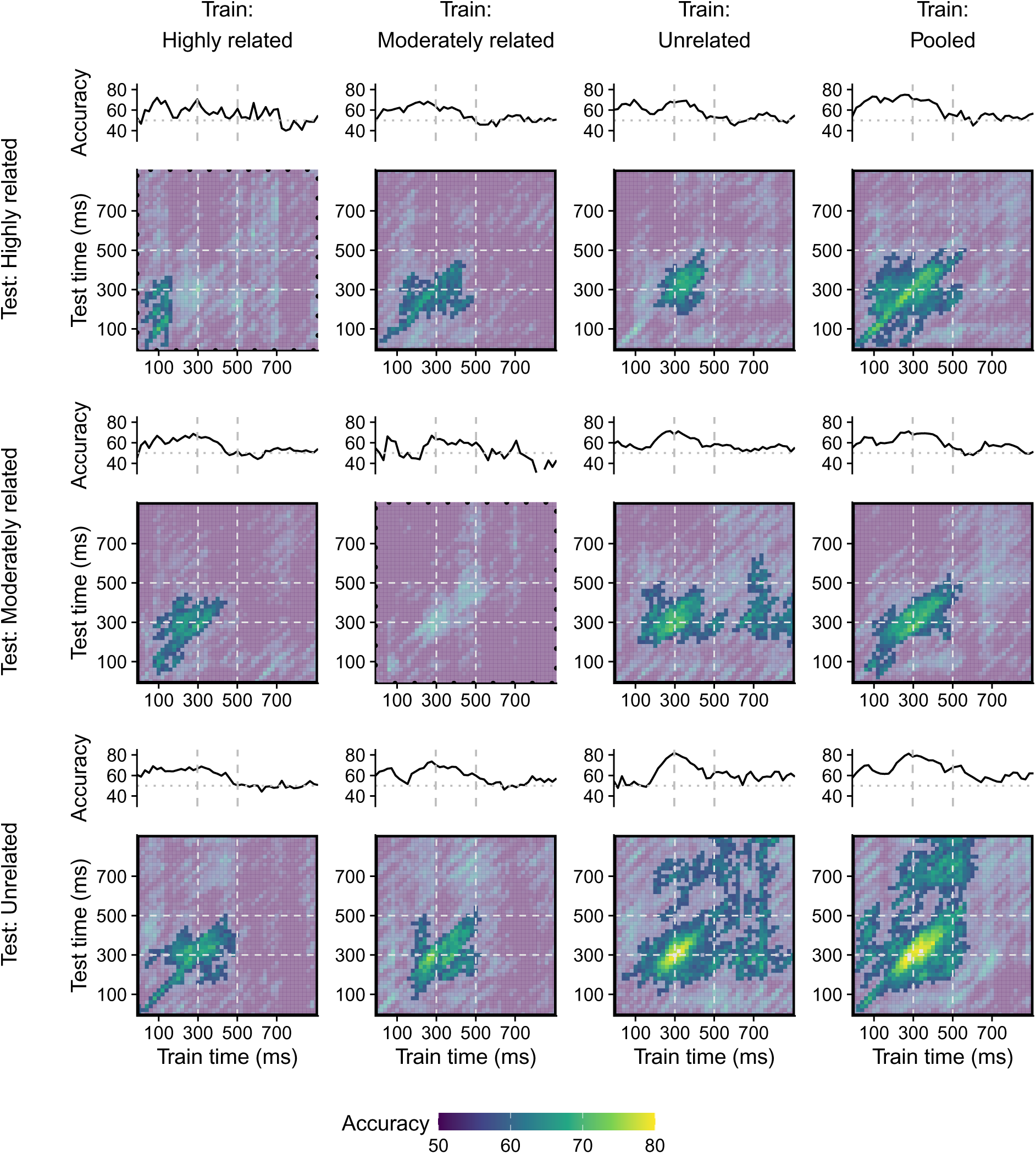
Temporal generalization maps for semantic decoding models. Here, models were trained on one 100-ms time window and tested on another 100-ms time window. Training and testing windows were either from the same or different relatedness levels. The color corresponds to the leave-two-out accuracies averaged over all targets. Clusters with associated *p*-values lower than .05 are highlighted. Dotted black borders indicates non-significance after FDR correction. Above each map are time-courses of decoding accuracy corresponding to the diagonals of the temporal generalization maps, which represent models that are trained and tested at the same time points.

**Figure 7:**
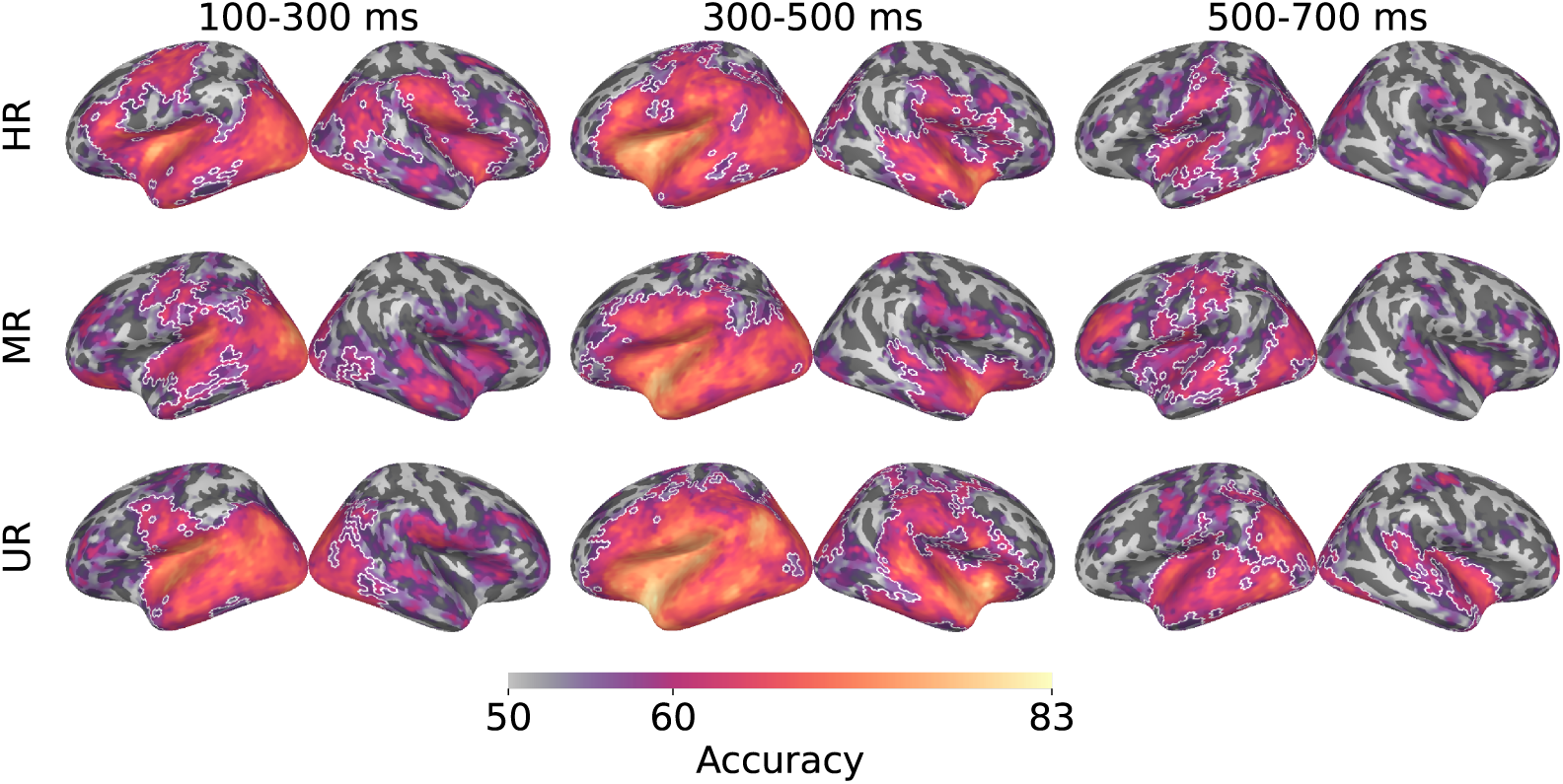
Decoding accuracy maps of cortical-level searchlight decoding across relatedness level and time window. White borders indicate clusters with associated *p*-values < 0.05. HR = highly related, MR = moderately related, UR = unrelated.

What occurs after the N400 window is notable. Decodability rapidly dropped relative to the previous windows, with only unrelated targets being decodable above chance (Figure 5, right). Although searchlight decoding indicated that some regions still presented context-invariant information for highly related and moderately related targets (Figure 7), the time-resolved decoding and temporal generalization maps showed a fast decline in decoding accuracy and generalization after around 500 ms (Figure 6). Accordingly, soon after the N400 peak, neural responses no longer seem to contain information that can be mapped to context-invariant word2vec embeddings. This suggests that the N400 window may index the last stage in the processing time-course when the stimulus is represented in a manner independent of the semantic context.

While the results indicate that representations of the target words were encoded in the neural response irrespective of contextual support, the differences between relatedness levels have implications for the training of decoders. Words unrelated to their preceding primes elicited higher N400 and provided the best training examples to learn the mapping between neural responses and semantic vectors. This finding can inform the development and training of future decoding models: focusing on examples that elicit high N400 responses may be a promising path towards creating an accurate and generalizable context-invariant semantic decoder.

Our study bridges two lines of research: classic evoked-response measures and modern representational analyses. These methods provide complementary information about semantic processing that cannot be inferred from either one alone. However, it is important to address the limitations. First, the decoding analyses were based on item-level grand-averaged MEG data. This approach improves signal-to-noise ratio and is valuable for detecting stable semantic structure, but it limits conclusions about whether comparable semantic information can be decoded at the single-trial level or within individual participants. Second, responses to written words involve multiple partially overlapping processing stages, including visual, orthographic, lexical, and semantic processing. Although the mapping to context-invariant semantic vectors supports a semantic interpretation of the decoding results, it may not fully separate semantic information from correlated perceptual processes. Third, due to the experimental design, we did not perform decoding analyses for the prime words, or for targets before the target word onset. However, future work could endeavor to track decodability of both primes and targets throughout a trial, building on studies that looked at multi-word contexts and predictive pre-activation (Dikker and Pylkkanen, 2013; Fyshe et al., 2019; Eisenhauer et al., 2022), potentially utilizing semantic embeddings that encode context such as BERT (Devlin et al., 2019) or GPT-2 (Radford et al., 2019).

In conclusion, our results bridge the classical paradigm of stimulus-evoked modulation of local neural activity and the more recent neural decoding approach. We elucidated the time course of semantic information as captured by distributed patterns of neural activity, with reference to the well-established N400 response.

## 5. Acknowledgements

We thank Kasper Rantamaki for assistance with stimuli generation scripts, and Tuomas Tolvanen and Mia Illman for technical support during data collection. We additionally thank Jenna Kanerva and Filip Ginter at the University of Turku for development of the Finnish language word2vec model. We acknowledge the computational resources provided by the Aalto Science-IT project.

Financial support for this project was provided by the Finnish Cultural Foundation (#00230334 and #00240405 to GG and #75242391 to AS), the Finnish Foundation for Technology Promotion (#11215 to GG), the Research Council of Finland (#346585 and #343385 to MvV and #355407 to RS) and the Sigrid Juśelius Foundation (to RS).

## 6. Declarations

The authors declare no conflicts of interest.

## Appendix A Stimuli details

**Figure 8:**
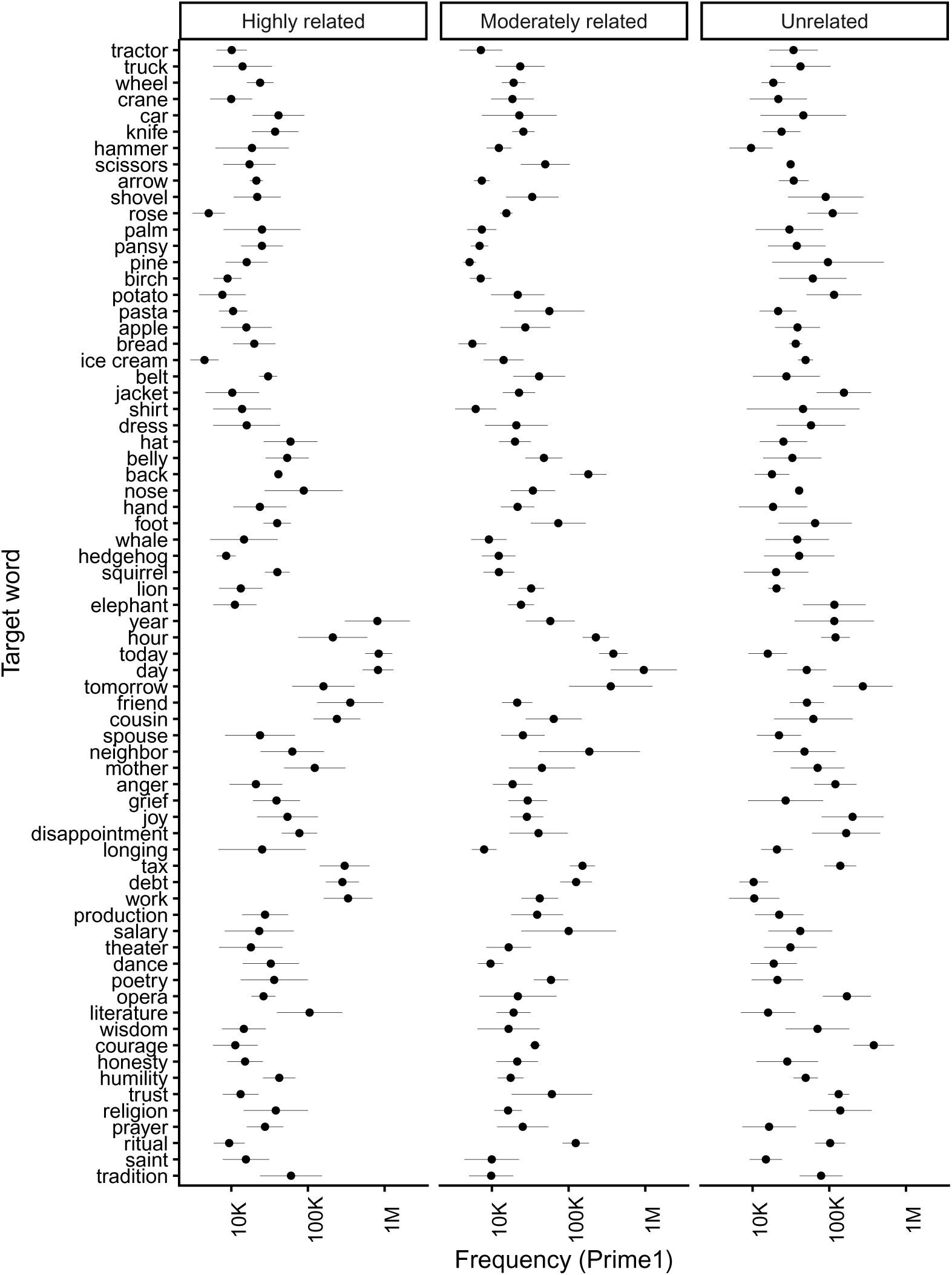
Word frequency (mean and standard error) of prime 1 words for each target (translated into English) in the three conditions.

**Figure 9:**
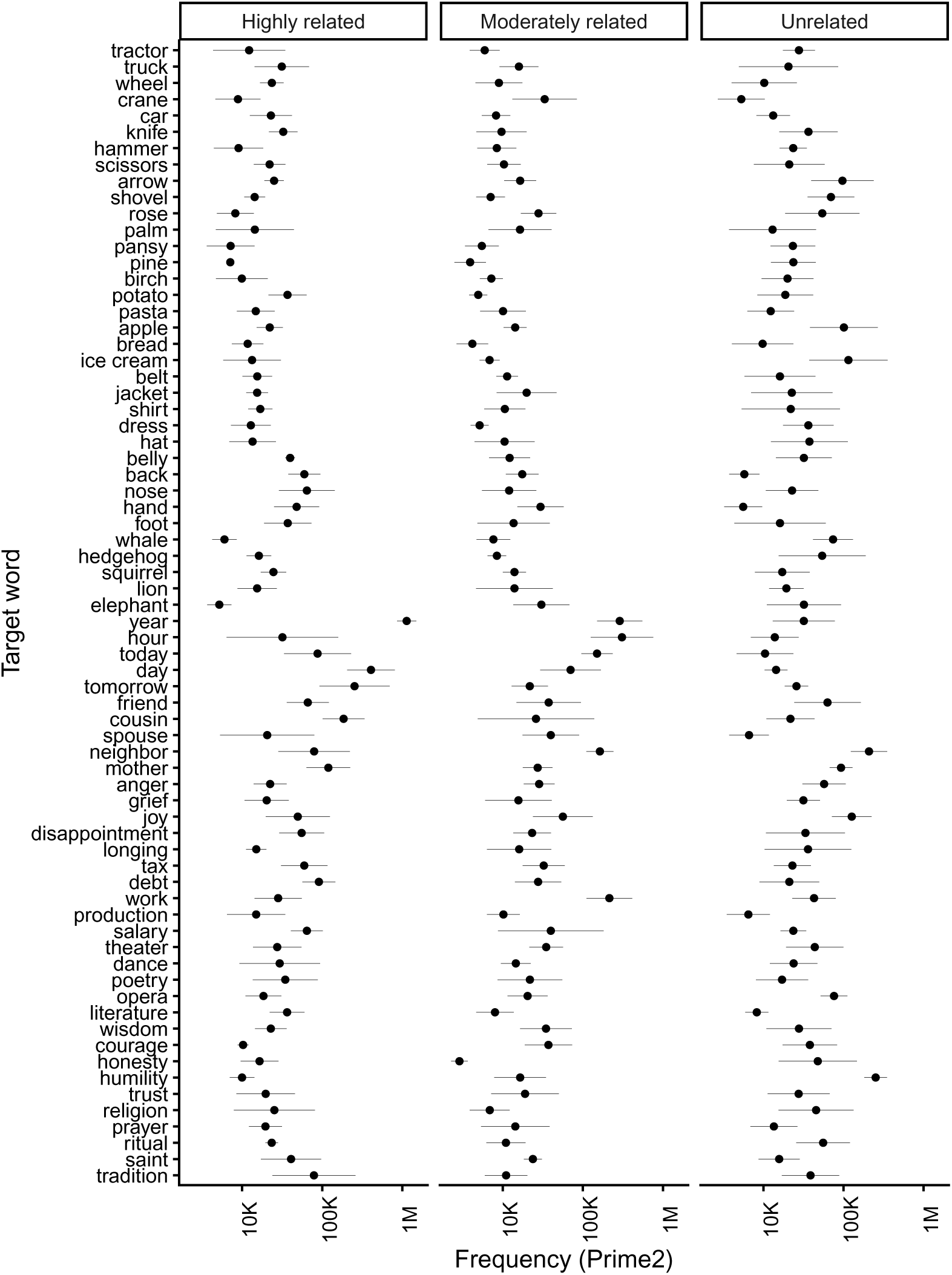
Word frequency (mean and standard error) of prime 2 words for each target (translated into English) in the three conditions.

**Figure 10:**
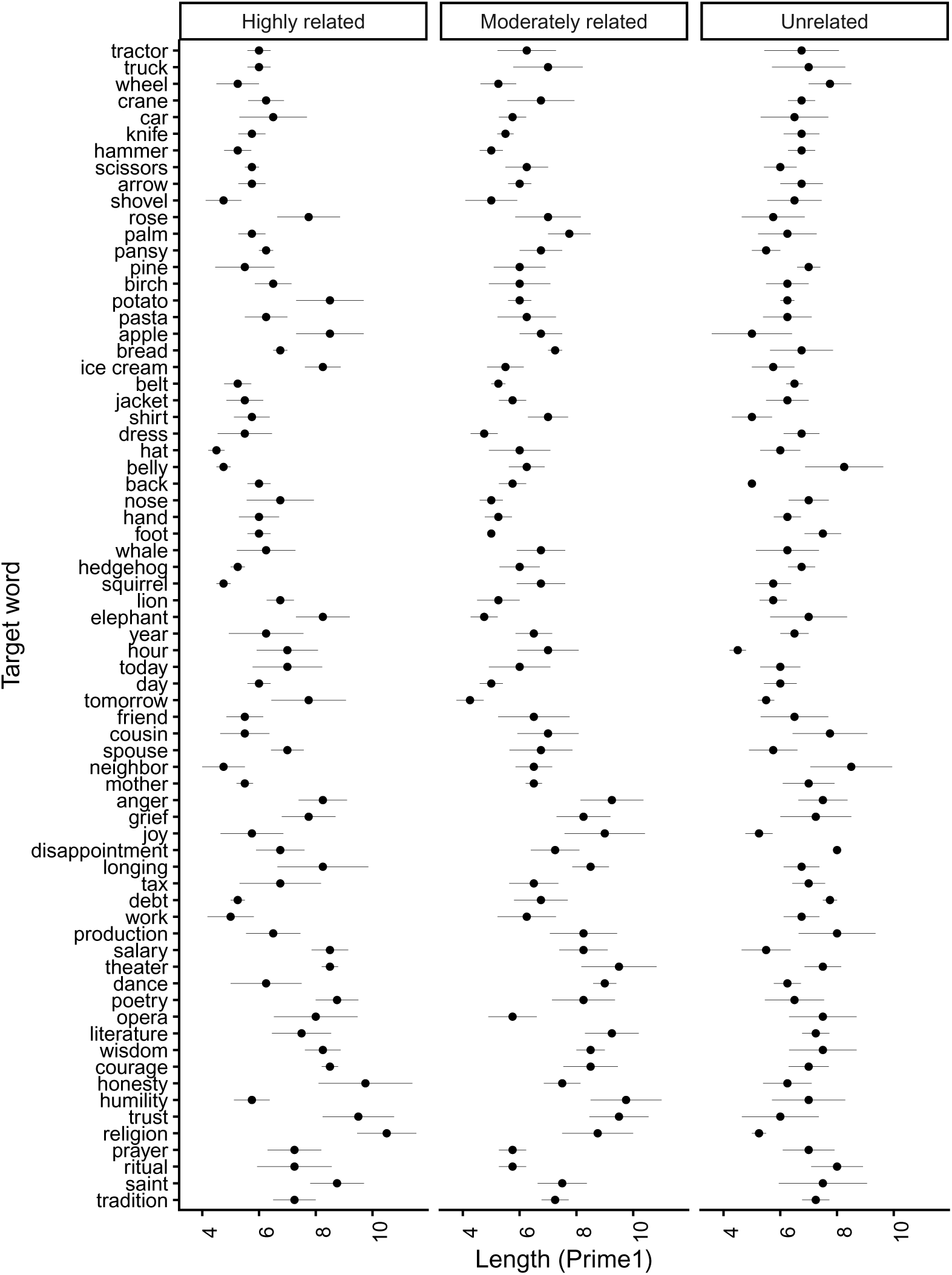
Word length (mean and standard error) of prime 1 words for each target (translated into English) in the three conditions.

**Figure 11:**
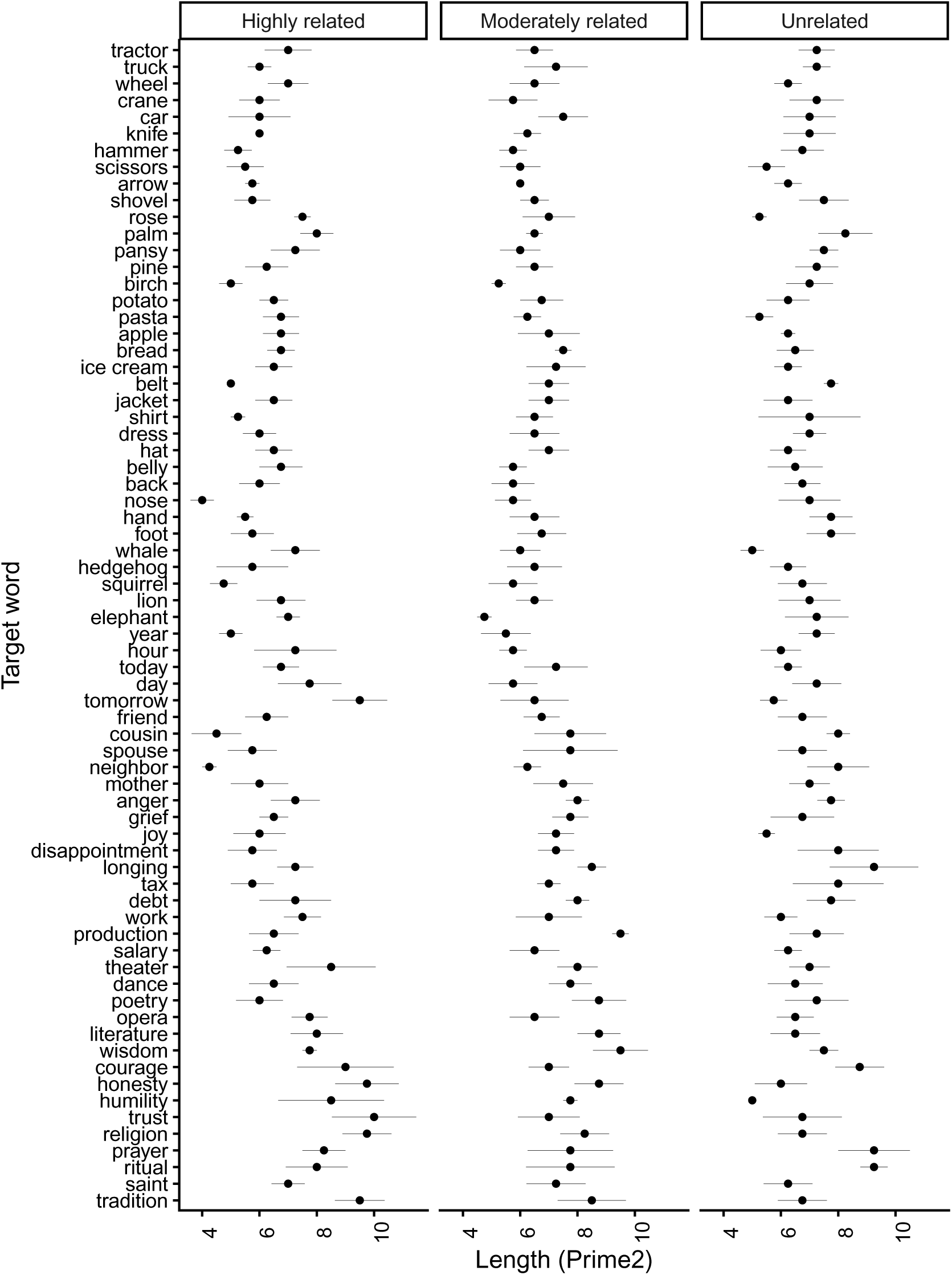
Word length (mean and standard error) of prime 2 words for each target (translated into English) in the three conditions.

## Notes

https://github.com/AaltoImagingLanguage/ghazaryan2026

### Competing Interest Statement

The authors have declared no competing interest.

### Summary of Updates

Minor updates to manuscript text and figures

